# Immune-evasion of *KRAS*-mutant lung adenocarcinoma mediated by cAMP response element-binding protein

**DOI:** 10.1101/2021.06.19.449094

**Authors:** Georgia A. Giotopoulou, Giannoula Ntaliarda, Antonia Marazioti, Ioannis Lilis, Foteini Kalogianni, Evanthia Tourkochristou, Nikolitsa Spiropoulou, Ioanna Giopanou, Magda Spella, Marianthi Iliopoulou, Aigli Korfiati, Theofilos Mantamadiotis, Christian Rosero, Torsten Goldmann, Sebastian Marwitz, Georgios T. Stathopoulos

**Author notes:** **Corresponding author:** Georgios T. Stathopoulos, MD PhD; Comprehensive Pneumology Center, Max-Lebsche-Platz 31, 1. OG, 81377 Munich, Germany. Equal first authors (Georgia A. Giotopoulou, Giannoula Ntaliarda, Antonia Marazioti, Ioannis Lilis).

## Abstract

cAMP response element-binding protein (CREB) mediates proliferative and inflammatory gene transcription in neurodegeneration and cancer, but its role in malignant immune-evasion of lung adenocarcinoma (LUAD) is unknown. We show that human LUAD of smokers are frequently altered along the CREB pathway and we employ mouse models to discover that *KRAS*-mutant LUAD co-opt CREB to evade immune rejection by tumoricidal neutrophils. For this, *KRAS*- driven CREB activation suppresses CXC-chemokine expression and prevents recruitment of CXCR1+ neutrophils. CREB1 is shown to be pro-tumorigenic in five different LUAD models, a function that is dependent on host CXCR1. Pharmacologic CREB blockade prevents tumor growth and restores neutrophil recruitment only when initiated before immune-evasion of *KRAS*-mutant LUAD. CREB and CXCR1 expression in human LUAD are compartmentalized to tumor and stromal cells, respectively, while CREB-regulated genes and neutrophils impact survival. In summary, CREB-mediated immune evasion of *KRAS*-mutant LUAD relies on signaling to neutrophil CXCR1 and is actionable.

---

Lung adenocarcinomas (LUAD) carrying mutations in the *KRAS* proto-oncogene GTPase are still clinically undruggable (1). In addition to its molecular structure that lacks a deep pocket (2), the mutant KRAS oncoprotein activates multiple intracellular signaling cascades and initiates inflammatory interactions with the host immunity and vasculature, rendering its inhibition troublesome (2-5). However, targeting downstream transcription factors that are addicted to and mediate oncogene effects presents a viable alternative strategy to inhibit the ability of mutant cells to survive and proliferate (6, 7). For example, inhibition of kinases that activate the NF-κB, WNT, and Myc pathways have been shown to be effective against *EGFR*- and *KRAS*-mutant LUAD and to circumvent resistance to targeted therapies (8-12).

Upon its activation via phosphorylation, the nuclear transcription factor cAMP response element-binding protein (CREB; encoded by the human/murine *CREB1/Creb1* genes) is a regulator of key cellular processes such as survival, growth, differentiation, and inflammatory signaling (13-16). In the lungs, CREB is an essential survival factor for the normal respiratory epithelium during pulmonary development and adult lung homeostasis (17). Since LUAD of smokers arise from the respiratory epithelium (18) and CREB signaling has been documented in such tumors (19), the transcription factor is ideally positioned as a candidate LUAD promoter that could mediate sustained survival and proliferation of tumor-initiated epithelial cells. However, its role in LUAD has not been assessed *in vivo*.

Neutrophils that express C-X-C-motif chemokine receptor 1 (CXCR1) and possess antitumor functions have been identified in pre-metastatic human LUAD (20, 21) and have also been recently documented in single-cell transcriptomic screens of human and mouse LUAD (22). Although experimental evidence for pro-tumor neutrophil functions in multiple cancer types is ample (23, 24), functional proof for the existence of anti-neoplastic neutrophil functions in mouse models of pre-metastatic LUAD do not exist.

We identified frequent alterations of the CREB pathway in human LUAD of smokers. To functionally interrogate their role, we used conditional deletion, forced overexpression, and pharmacologic inhibition of CREB in different mouse and cellular models of early-stage *KRAS*-mutant LUAD to identify a marked tumor-promoting role of this transcription factor. Importantly, CREB effects were druggable and were delivered through a novel mechanism of immune-evasion mediated via paracrine suppressor signaling to host neutrophils.

## RESULTS

### CREB signaling drives *KRAS*-mutant LUAD

A query of mutations, copy number alterations, fusions, and mRNA expression in The Cancer Genome Atlas (TCGA) LUAD dataset (25) revealed frequent alterations of the CREB signaling pathway (composed of the *NGF, NRN1, NTRK1/2, CREB1*, and *CREBBP* genes) in 41% of patients (**Figs. 1A-C**). CREB pathway alterations were associated with increased tumor stage, mutation count, and C>A transversions, suggesting linkage with smoking, while most *CREB1* gene alterations consisted of mRNA overexpression, both in *KRAS*-altered and normal patients (**Figs. 1D-F**).

**Fig. 1.**
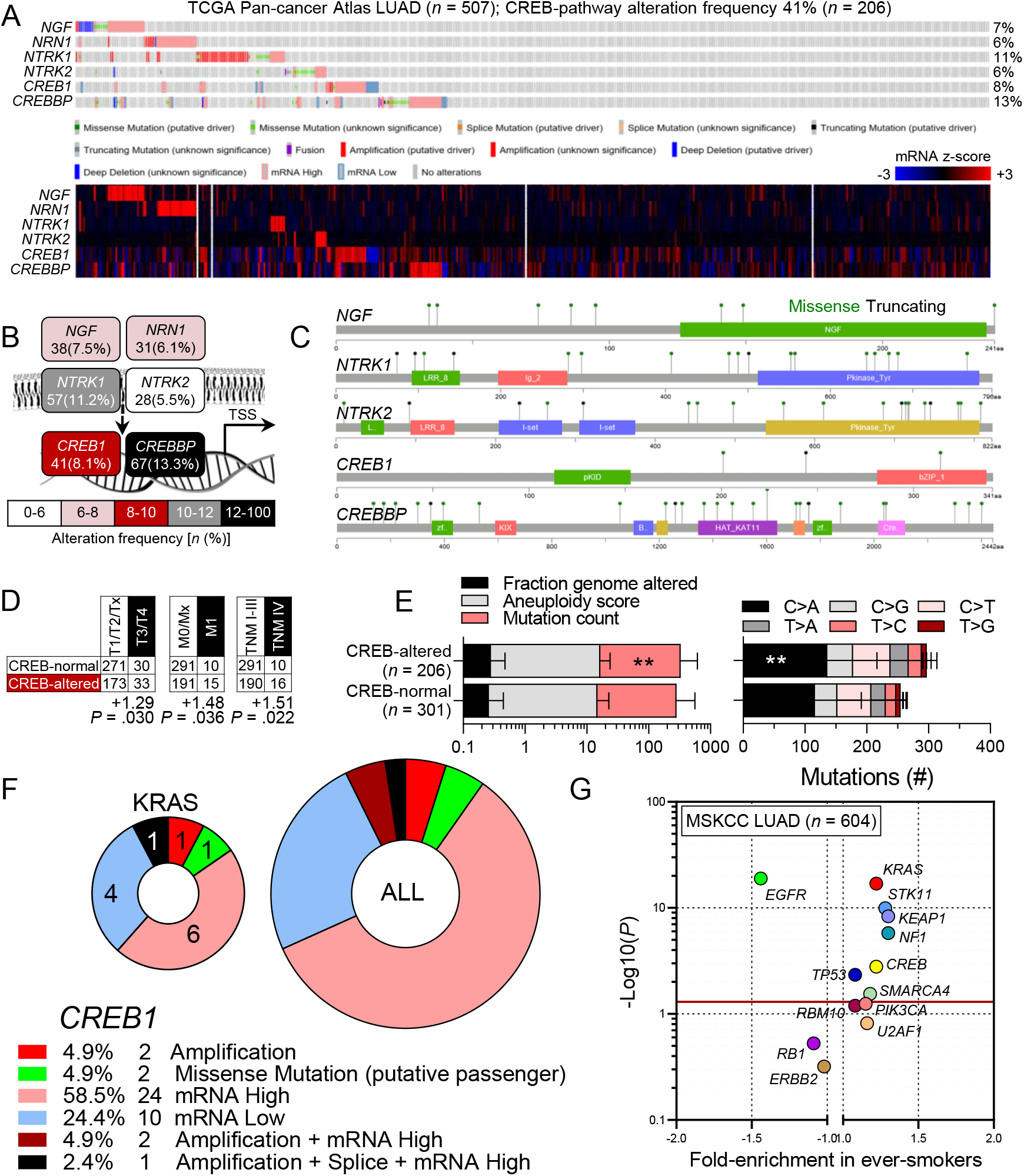
CREB pathway alterations in human lung adenocarcinoma (LUAD). Data from the cancer genome atlas (TCGA; *n* = 507; A-F) and Memorial Sloan Kettering Cancer Center (MSKCC; *n* = 604; G) datasets (https://www.cbioportal.org/; TCGA link: https://bit.ly/3vfWDYn; MSKCC link: https://bit.ly/3pMBJz9). **(A)** Mutation plot (top) and gene expression heatmap (bottom) with gene names, alteration frequencies, legend, and color key. Each column represents one patient. **(B)** Pathway schematic with color-coded alteration frequencies. **(C)** Mutation lollipop plots. Note the presence of over/under-expression and the absence of mutations in *NRN1*. **(D)** Cross-tabulations of tumor stage by CREB status shown as patient numbers with hypergeometric test enrichment and probability (*P*). **(E)** Data summary of genomic alterations (left) and signatures (right). **: *P* < 0.01 compared with CREB-normal patients, 2-way ANOVA, Sidak’s post-test. **(F)** *CREB1* alteration types in all (big pie chart) and in *KRAS*-altered (small pie chart) patients with legend, alteration frequencies, and patient numbers. Table refers to all patients. Numbers in small pie chart refer to *KRAS*-altered patients. **(G)** Hypergeometric test fold-enrichment versus *P* for CREB pathway mutations versus ever smoking in the MSKCC dataset. Red line represents *P* < 0.05 cut-off. Note the statistically significant co-segregation of CREB pathway mutations with *KRAS, STK11, KEAP1, NF1, TP53*, and *SMARCA4* mutations in ever smokers. Data shown as mean±SD, patient numbers (*n*), and/or percentages (%).

In the Memorial Sloan Kettering Cancer Center (MSKCC) dataset, CREB pathway alterations co-segregated with *KRAS, STK11, KEAP1, NF1, TP53*, and *SMARCA4* mutations in ever smokers (**Fig. 1G**). Hence, to study *Creb1* function in LUAD, we employed five different mouse models that feature *KRAS* mutations and/or *TP53* loss. First, in a model of early-stage *Kras*^Q61R/G12V^-mutant LUAD with six months latency (12, 18), wild-type (*Creb1*^WT/WT^) and conditional *Creb1*-deleted (*Creb1*^F/F^) mice (26) back-crossed > F12 onto the carcinogen-susceptible FVB strain received 1 g/Kg i.p. urethane (ethyl carbamate, a tobacco carcinogen). Two weeks prior to urethane administration, mice received 5 × 10^9^ PFU intratracheal (i.t.) adenovirus-type-5 encoding CRE recombinase (Ad-CRE) that results in 90% recombination of respiratory epithelial cells within 2 weeks (12). Second, Scgb1a1.CRE mice that express CRE in club cells (12, 18) were intercrossed with *Creb1*^WT/WT^ and *Creb1*^F/F^ mice (all > F12 FVB), and the offspring received urethane as above. Third, LSL.*KRAS*^G12D^mice (C57BL/6 strain) that carry a conditional loxP-STOP-loxP.*KRAS*^G12D^ transgene (12, 18, 22) were intercrossed with *Creb1*^WT/WT^ and *Creb1*^F/F^ mice, and their offspring received 5 × 10^8^ PFU i.t. Ad-CRE, resulting in sporadic LUAD with *KRAS*^G12D^ mutations after four months. Fourth, a primary LUAD cell line was established by culture of a urethane-induced LUAD from a non-Ad-CRE-treated *Creb1*^F/F^ mouse as described previously (27), providing an *in vitro* model for *Creb1*-deletion in the cellular context of *Kras*^G12V^-mutant LUAD (**Figs. S1A-E**). As a fifth approach, benign human HEK293T epithelial cells were stably transfected with plasmid encoding murine *Kras*^G12C^ (p*Kras*^G12C^) plus plasmids encoding *Photinus pyralis* luciferase (p*LUC*) or murine *Creb1* (p*Creb1*) and 3 × 10^6^ cells were injected s.c. into the bilateral flank dermis of *NOD/SCID* mice. In all five models, the expression of nuclear phosphorylated activated CREB (P-CREB) was increased compared to its sporadic expression in the naïve pulmonary epithelium and to other mouse (Lewis lung carcinoma, LLC) and human (A549) LUAD cell lines, and showed the highest immunoreactivity at the tumor-host interface (**Figs. S2A-C**). *CREB1* mRNA was also markedly overexpressed in published LUAD transcriptomes compared with normal lungs from never smokers (**Fig. S2D**). Importantly, conditional deletion of *Creb1* during LUAD development markedly decreased lung tumor burden in all five *in vivo* models examined, achieving 74% reduction in tumor burden overall (**Figs. 2A-E**). CREB signaling is required for the development of *KRAS*-mutant LUAD, since only 5/112 (4%) *Creb1*-competent tumors were detected in urethane-treated *Creb1*^F/F^ mice (**Figs S3A-D**) and none of the 507 TCGA LUAD patients had a *CREB1* deletion (**Fig. 1F**). CREB activation in primary LUAD cells was not driven by the coincident *Trp53* loss these cells feature (27), but by mutant *Kras* (**Figs. S4A-F**). These results present robust *in vivo* evidence indicating that CREB is required for *KRAS*-mutant LUAD in humans and mice.

**Fig. 2.**
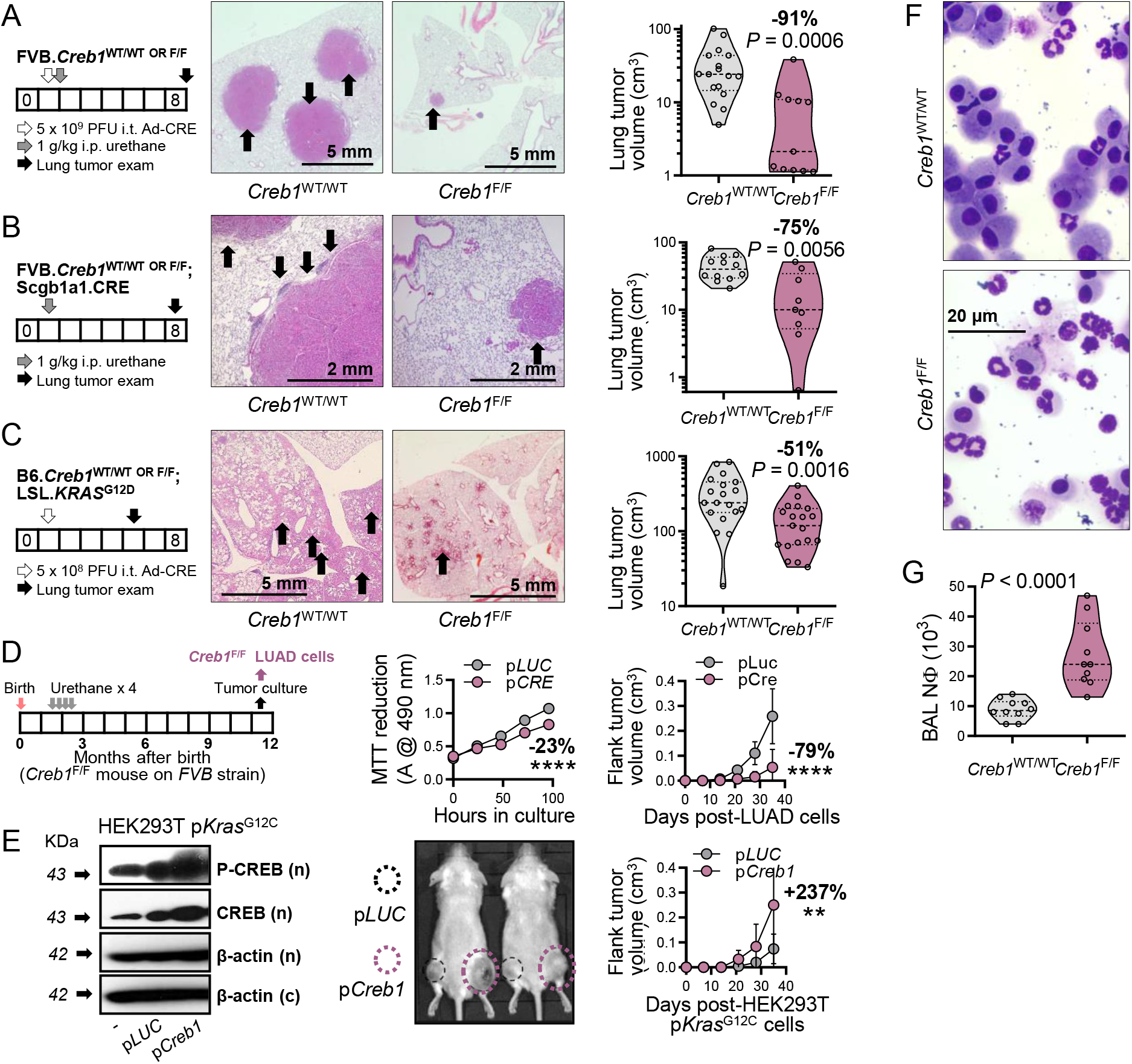
CREB is a LUAD promoter. **(A-C)** Experimental setup (schematics; boxes = months), representative H&E-stained lung sections (images), and pooled results (graphs) from *Creb1*^WT/WT^ (*n* = 17) and *Creb1*^F/F^ (*n* = 11) mice (FVB strain) at 6 months after i.t. 5 × 10^9^ PFU Ad-CRE followed by i.p. 1 g/Kg urethane 2 weeks later (A), from Scgb1a1.CRE;*Creb1*WT/WT (*n* = 12) and Scgb1a1.CRE;*Creb1*^F/F^ (*n* = 9) mice (FVB strain) at 6 months after i.p. 1 g/Kg urethane (B), and from LSL.*KRAS*^G12D^;*Creb1*^WT/WT^ (*n* = 19) and Scgb1a1.CRE;*Creb1*^F/F^ (*n* = 19) mice (C57BL/6 strain) at 4 months after i.t. 5 × 10^8^ PFU Ad-CRE. Arrows in images denote LUAD. **(D)** *Creb1*^F/F^ LUAD cells were derived from a LUAD induced by four weekly i.p. injections of 1 g/Kg urethane in a FVB *Creb1*^F/F^ mouse (schematic; boxes = months), were stably transfected with p*LUC* or p*CRE*, were validated, and were assessed for MTT reduction (*n* = 4/data-point) and tumor growth in FVB mice (p*LUC, n* = 12; p*CRE, n* = 10). **(E)** HEK293T cells stably transfected with p*Kras*^G12C^ plus p*LUC* or p*Creb1*, were validated (immunoblots; n, nuclear; c, cytoplasmic) and 3 × 10^6^ cells were injected s.c. into *NOD/SCID* mice(*n* = 6/group). (D, E) Shown are representative photographs and data summaries as mean (circles) and SD (bars). *P*, probability, two-way ANOVA. ** and ****: *P*< 0.01 and *P*< 0.0001, respectively, compared with p*LUC*, Bonferroni post-tests. **(F, G)** Representative May-Grünwald-Giemsa-stained cytocentrifugal specimens (F) and data summary (G; *n* = 10/group) of bronchoalveolar lavage (BAL) neutrophils (NΦ) from LUAD-bearing *Creb1*^WT/WT^ and *Creb1*^F/F^ mice. Data in (A-C, G) are shown as raw data points (circles), rotated kernel density plots (violins), medians (dashed lines), and interquartile ranges (dotted lines). *P*, probability, Mann-Whitney U-test.

### CREB prevents recruitment of tumoricidal neutrophils

Surprisingly, all LUAD-protected *Creb1*-deleted mice displayed increased numbers of CD45^+^CD11b^+^Gr1^+^CXCR1^+^ neutrophils in bronchoalveolar lavage (BAL) as determined by cytology and flow cytometry (**Figs. 2F, S5A, S5B**). Moreover, supernatants of bone marrow-derived neutrophils matured via 48-hour incubation with 20 ng/mL G-CSF (5) potentiated the anti-tumor activity of the KRAS inhibitor deltarasin (**Figs. S5C-E**). Microarray analysis of *Kras*^G12V^-mutant *Creb1*^F/F^primary LUAD cells stably infected with p*LUC* or with p*CRE* (GEO dataset GSE156513) identified 486 murine genes driven/suppressed by CREB signaling, hereafter called the CREB signature. Intertestingly, the CXCR1/2 ligands *Cxcl3, Cxcl5*, and *Ppbp* were among the top CREB-suppressed genes, and CREB deletion up-regulated inflammatory signaling pathways in LUAD cells (**Figs. S6A-E**). To this end, CREB-binding motifs (5’-TGACGTCA-3’; ENCODE identifier ENCFF576PUH) were found in the promoters of human CXC chemokine genes using public sequencing/ChIP data from the encyclopedia of DNA elements (ENCODE) (32) (**Fig. S7**). *Cxcr1*-deficient mice (33) displayed the opposite phenotype of *Creb1*-deficient animals exhibiting marked susceptibility to chemical and transplantable LUAD and showing impaired neutrophil recruitment to LUAD (**Figs. 3A-C**). Host *Cxcr1* deficiency abrogated the enhanced tumor growth provided by *Creb1* competence, mutant *Kras* in tumor cells was required for the tumor-prone phenotype of *Cxcr1*-deficient mice, and tumor-restricted *Creb1* deletion resulted in diminished P-CREB but enhanced CXCR1 tumor immunoreactivity (**Figs. 3D-G**). Collectively, these results indicate that, in *KRAS*-mutant LUAD, CREB activation silences chemokine signaling to prevent neutrophil-mediated tumor rejection.

**Fig. 3.**
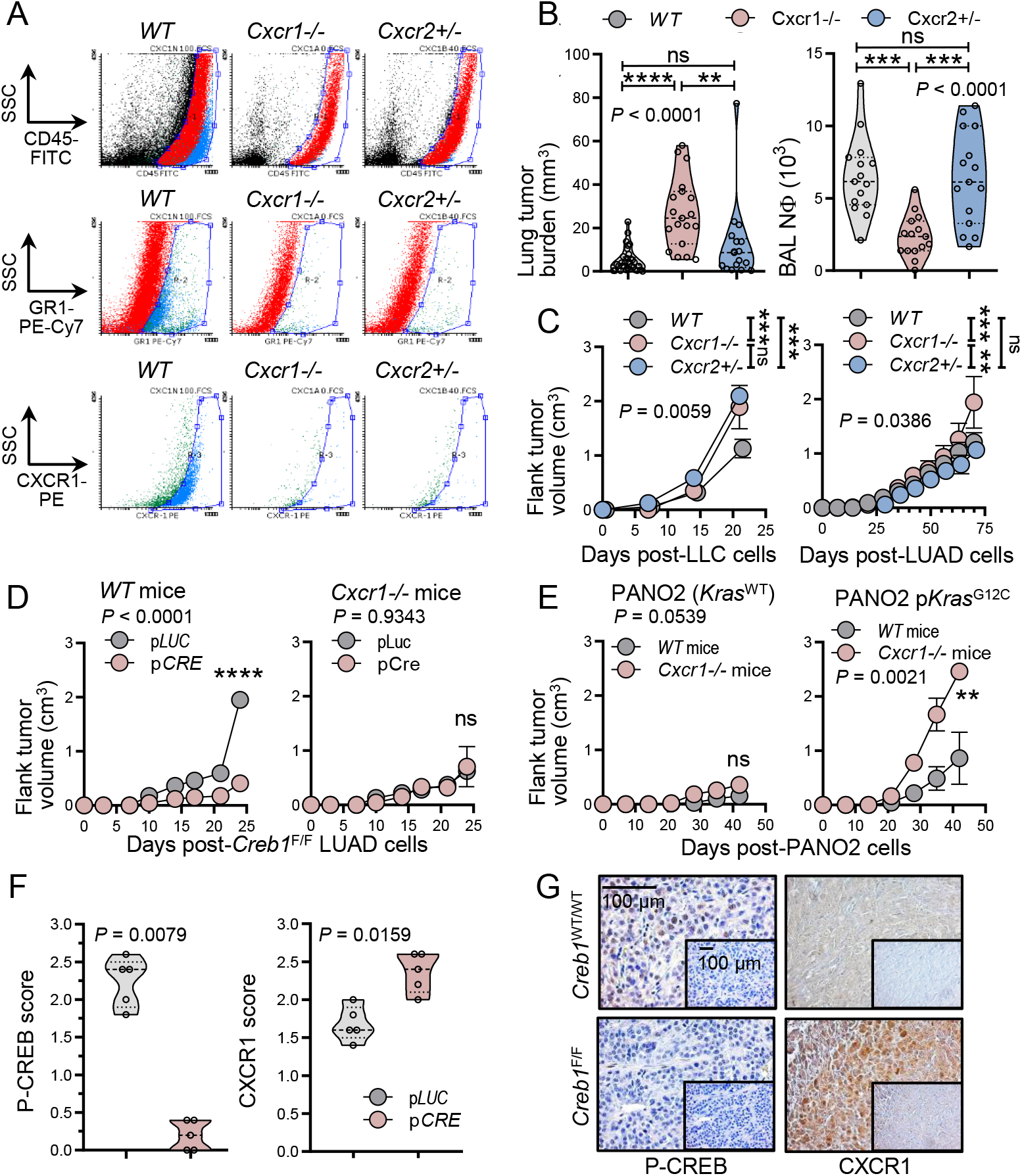
CREB signaling prevents neu1 trophil influx into the LUAD-affected lungs. CXCR1+ neutrophils facilitate immune surveillance. **(A)** Representative May-Grünwald-Giemsa-stained cytocentrifugal specimens of broncho0alveolar lavage (BAL) from LUAD-bearing *Creb1*^WT/WT^ and *Creb1*^F/F^ mice. **(B)** Data summary (*n* = 0105/gr1o0 u1p5)2o0 f 2c5ytology for BAL neutrophils (NΦ) shown as raw data points (circles), rotated kernel density plots (violins), medians (dashed lines), and interquartile ranges (dotted lines). *P*, probability, Mann-Whitney U-test.**(A)**Data summary of lung tumor burden (left) and CD45+CD11b+Gr1+ bronchoalveolar lavage (BAL) neutrophils (NΦ; right) of *WT, Cxcr1*^*-/-*^, and *Cxcr2*^*+/-*^ mice at 6 months post-1 g /Kg i.p. urethane shown as raw data points (circles), rotated kernel density plots (violins), medians (dashed lines), and interquartile ranges (dotted lines). Tumor burden: *n* = 32, 19, and 17, respectively, for *WT, Cxcr1*^*-/-*^, and *Cxcr2*^*+/-*^ mice. BAL NΦ: *n* = 15/group. *P*, probabilities, one-way ANOVA. ns, **, ***, and ****: *P* > 0.05, *P* < 0.01, *P* < 0.001, and *P* < 0.0001, respectively, for the indicated comparisons, Tukey’s post-tests.**(B)**Data summary of primary tumor volume after s.c. injection of 10^6^ LLC (left) and primary urethane-induced LUAD (right) cells into *WT, Cxcr1*^*-/-*^, and *Cxcr2*^*+/-*^ mice presented as mean (circles) and SD (bars). LLC cells: *n* = 14, 9, and 6, respectively. LUAD cells: *n* = 8, 4, and 4, respectively. *P*, probabilities, two-way ANOVA. ns, **, and ***: *P* > 0.05, *P* < 0.01, and *P* < 0.001, respectively, Bonferroni post-tests. **(C, D)**Data summary of primary tumor volume after s.c. injection of 10^6^ *Creb1*^F/F^ primaryLUAD cells stably expressing p*LUC* or p*CRE* into *WT* (left) and *Cxcr1*^*-/-*^ (right) mice (C) and of 10^6^ pancreatic adenocarcinoma (PANO2) cells stably expressing p*C* (left) or p*Kras*^G12C^ (right) into *WT* and *Cxcr1*^*-/-*^ mice (D) presented as mean (circles) and SD (bars). *n* = 6/data point. *P*, probabilities, two-way ANOVA. ns, **, and ****: *P* > 0.05, *P* < 0.01, and *P* < 0.0001, respectively, Bonferroni post-tests. **(E, F)** Data summary (left; *n* = 5/group) and representative CXCR1 (brown) immunohistochemistry images (blue: hematoxylin) of tumor sections from experiment in C (left) shown as raw data points (circles), rotated kernel density plots (violins), medians (dashed lines), and interquartile ranges (dotted lines). *P*, probability, Mann-Whitney U-test.

### CREB blockade prevents immune evasion of *KRAS*-mutant LUAD and restores neutrophils, a hallmark of early-stage disease

To block CREB signaling, FVB mice bearing s.c. *Kras*^G12V^ LUAD received i.p. CREB inhibitors before or after tumor establishment. While ICG-001, a CBP/β-catenin interaction inhibitor used here as internal control (36) was ineffective, KG501 (Naphthol-AS-E-phosphate), a specific CREB1-CBP interaction inhibitor (37), prevented CREB activation, restored LUAD neutrophils, and blocked tumor development only when administered early after tumor establishment (Figs. 4A-C). More neutrophils infiltrated the stroma compared with tumor areas of 195 patients with all histologic subtypes of lung cancer (**Figs. 4D-E**). However, the abundance of stroma-infiltrating neutrophils was associated with improved survival specifically of 101 patients with LUAD (**Figs. 4F-G**). In addition, the CREB transcriptome signature was enriched in human smokers’ LUAD from the BATTLE study (GSE43458) and predicted poor survival of LUAD in the KMploit dataset (https://kmplot.com/) (**Figs. S8A-D**). An inverse correlation between tumor cell CREB and stromal cell CXCR1 protein expression was also striking in 42 patients from the Human Protein Atlas (http://www.proteinatlas.org), further supporting our hypothesis (**Figs. S9A-C**). We conclude that CREB mediates immune evasion of *KRAS*- mutant LUAD by inhibiting chemokine secretion and by preventing the recruitment of tumoricidal neutrophils (**Fig. S9D**).

**Fig. 4.**
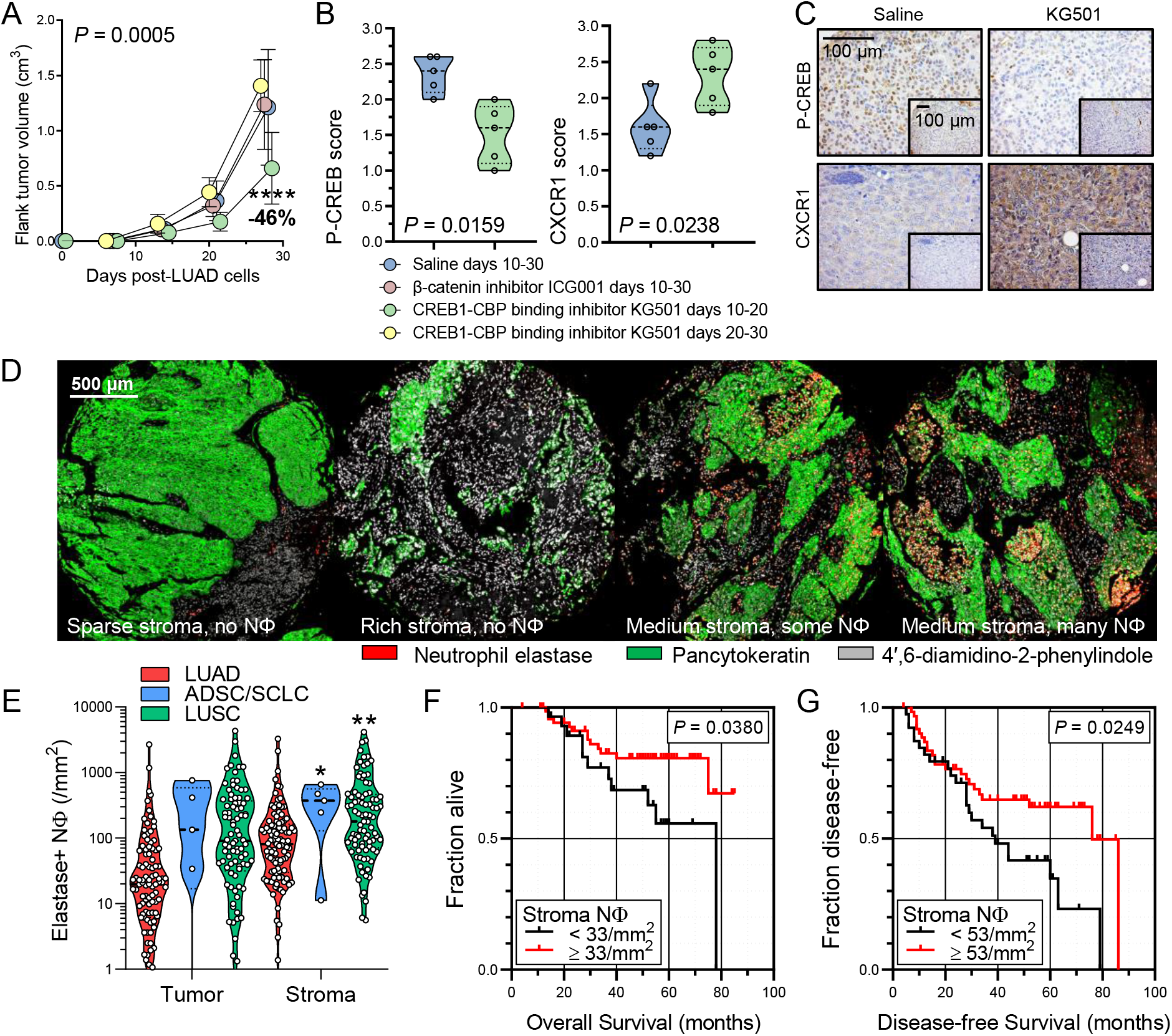
CREB-mediated immune evasion of *KRAS*-mutant LUAD is druggable and likely occurs in human LUAD. **(A)** Data summary of primary tumor volume after injection of 10^6^ s.c. primary LUAD cells into FVB mice followed by treatment with daily saline during experimental days 0-30 (*n* = 16), the β-catenin inhibitor ICG-001 during experimental days 10-30 (*n* = 9), the CREB-CREB binding protein (CBP) inhibitor KG501 during experimental days 10-20 (*n* = 6), or KG501 during experimental days 20-30 (*n* = 6) post-LUAD cells presented as mean (circles) and SD (bars). *P*, probability, two-way ANOVA. ****: *P* < 0.0001, Bonferroni post-test. Only significant differences are indicated. **(B, C)** Data summary (B; *n* = 5/group) and representative P-CREB and CXCR1 (brown) immunohistochemistry (blue: hematoxylin) images (C; inlays, isotype controls) of tumor sections from experiment in (A) shown as raw data points (circles), rotated kernel density plots (violins), medians (dashed lines), and interquartile ranges (dotted lines). *P*, probability, Mann-Whitney U-test. **(D-G)** Representative multi-color immunohistochemistry images (D), data summary (E), and Kaplan-Meier survival plots (F, G) of human lung tumors (LUAD, *n* = 101; adenosquamous, ADSC, *n* = 3, small cell lung cancer, SCLC, *n* = 2; squamous cell lung carcinoma, LUSC, *n* = 89). Data in (E) are shown as raw data points (circles), rotated kernel density plots (violins), medians (dashed lines), and interquartile ranges (dotted lines). * and **, P < 0.05 and P < 0.01, respectively, for comparison with tumor, 2-way ANOVA with Sidak’s post-tests. Data in (F, G) are shown as Kaplan-Meier survival estimates (lines), censored observations (line marks), and log-rank probabilities (*P*).

## DISCUSSION

Here we describe the tumor-promoting functions of CREB in *KRAS*-mutant LUAD. We query six human cohorts and employ five different mouse models to discover that *KRAS*- mutant LUAD co-opt CREB in order to evade immune rejection by tumoricidal neutrophils. To this end, CREB activation prevents KRAS signaling to neutrophil CXCR1 via cognate CXCL chemokines thus prohibiting the recruitment of neutrophils. These granulocytes are shown to express CD45, CD11B, GR1, CXCR1, and neutrophil elastase and to directly synergize with a KRAS inhibitor to specifically kill *KRAS*-mutant tumor cells in vitro. CREB expression and its molecular signature appear to be enriched in smokers’ tumors, which also feature frequent *KRAS* mutations (25), are inversely correlated with CXCR1 expression, and predict poor survival. The KRAS-CREB-CXCR1 signaling loop is actionable using an inhibitor of the interaction between CREB-CREB binding protein (CBP) lending clinical implications to the data.

This is the first *in vivo* study of CREB effects in LUAD. Together with published work in other tumor types, the data support a potential pan-cancer role for the transcription factor (28, 29). The findings are mechanistically intriguing and present a leap forward over existing evidence from observational and *in vitro* studies (19, 30, 31). The results are also clinically important because *KRAS*-mutant LUAD is frequent and lethal and only recently became actionable with regard to G12C mutations (https://www.fda.gov/drugs/resources-information-approved-drugs/fda-grants-accelerated-approval-sotorasib-kras-g12c-mutated-nsclc). A novel mechanism is shown, which is functionally related to the identification of antigen-presenting neutrophils in human early-stage lung tumors (20, 21), and adds to existing insights of CREB signaling in adaptive immunity (34, 35). While we directly show their tumoricidal activity, CD45^+^CD11B^+^GR1^+^CXCR1^+^NE^+^ neutrophils remain to be further characterized at single cell resolution and to be functionally annotated for clinical significance in human disease (20-22).

In conclusion, CREB is activated in *KRAS*-mutant, but likely also in other molecular varieties of smokers’ LUAD with *TP53* and *STK11* mutations (19), and mediates immune evasion of these tumors from tumoricidal neutrophils, a mechanism that can be blocked by CREB-CPB interaction inhibition.

## METHODS

### Study approval

All animal experiments were prospectively approved by the Veterinary Administration of the Prefecture of Western Greece (#118021/579/30.04.2014) and were conducted according to Directive 2010/63/EU (http://eurlex.europa.eu/legal-content/EN/TXT/?uri=CELEX%3A32010L0063). The human study was approved *a priori* by the Ethics Committee of the University of Lübeck, Germany (#AZ 21-166), abided by the Declaration of Helsinki, and all patients gave written informed consent.

### TCGA analyses

TCGA pan-cancer atlas and MSKCC LUAD data were downloaded from https://www.cbioportal.org/ (38, 39) and were analyzed manually using Prism v8.0 (GraphPad, La Jolla, CA, RRID:SCR_002798).

### Reagents

Adenoviruses were from the Vector Development Lab, Baylor College of Medicine (Houston, TX) and 3-(4,5-dimethylthiazol-2-yl)-2,5-diphenyltetrazolium bromide (MTT) assay and urethane (ethyl carbamate, EC; CAS# 51-79-6) from Sigma (St. Louis, MO). Primer sets were *Kras*, CCATTTCGGACCCGGAG- CTTTAGTCTCTTCCACAGGCA; *Trp53*, CGCCGACCTATCCTTACCAT- TTCTTCTTCTGTACGGCGGT; *Creb1*, TCATGGTCGTTTTTATGT- AAAAGGGAAACAGGAAATGC-GGCATTGACACATATGCATAAAAC; *Gusb*, CTACTTGAAGATGGTGATCGCTC-ACAGATCACATCCACATACGG; *Mela*, CTGGACTCACTCCCTGTATCTC-TGGCTCGTCTTTCAAATTGGT; *Crebbp*, CAGTGAATCGCATGCAGGTTT-GAACTGAGGCCATGCTGTTC; *Ppbp*, GTACAGGCCAGGAGTTCACT-GACGATTCTCTTGACGCCAG; *Scgb1a1*, ATCACTGTGGTCATGCTGTCC-GCTTCAGGGATGCCACATAAC; *Mycoplasma Spp*, GGGAGCAAACAGGATTAGATACCCT-TGCACCATCTGTCACTCTGTTAACCTC, and *Cxcl5*, CTGCGTTGTGTTTGCTTAACC-TTCAGTTTAGCTATGACTTCCACC. Antibodies and dilutions were anti-P-CREB/CREB (Cell signaling, Danvers, MA; 9198S/9197S, RRID:AB_2561044/AB_331277, 1:1000), anti-histone 3 (Santa Cruz, Sta. Clara, CA; sc-518011, RRID:AB_2861152, 1:500), anti-β-actin (Santa Cruz, Sta. Clara, CA; sc-47778, RRID:AB_626632, 1:500), and anti-CXCR1 (Novus Biologicals; NBP2-16043, RRID:AB_2891014, 1:200), goat anti-rabbit (Southern Biotech; 4030-05, RRID:AB_2687483, 1:10000). Deltarasin (CAS #1440898-82-7) was from Tocris Bio-Techne (#5424; Wiesbaden-Nordenstadt, Germany).

### Experimental mice

C57BL/6J (C57BL/6; #000664, RRID:IMSR_JAX:000664), FVB/NJ (FVB; #001800, RRID:IMSR_JAX:001800), B6.129S4-*Kras*^tm4Tyj^/J (LSL.*KRAS*^G12D^ #008179, RRID:IMSR_JAX:008179), B6.129P2-*Cxcr1*^tm1Dgen^/J (*Cxcr1*^*-/-*^#005820, RRID:IMSR_JAX:005820), B6.129S2(C)-*Cxcr2*^tm1Mwm^/J (*Cxcr2*^*+/-*^#006848, RRID:IMSR_JAX:006848), NOD.Cg-*Prkdc*^*scid*^/J (*NOD/SCID*, #001303, RRID:IMSR_JAX:001303) were from Jackson Laboratories (Bar Harbor, MN). B6;CBATg(Scgb1a1-cre)1Vart/Flmg (Scgb1a1.CRE; European Mouse Mutant Archive #EM:04965, RRID:IMSR_EM:04965) are described elsewhere (40) and Creb1^tm3Gsc^ (*Creb1*^F/F^, Mouse Genome Informatics #MGI:2181395, RRID:IMSR_EM:02151) mice were donated by their founder (26). All mice were bred at the Center for Animal Models of Disease of the Department of Physiology at the Faculty of Medicine of University of Patras, Greece. All animal experiments were prospectively approved by the Veterinary Administration of the Prefecture of Western Greece (#118021/579/30.04.2014) and were conducted according to Directive 2010/63/EU (http://eurlex.europa.eu/legal-content/EN/TXT/?uri=CELEX%3A32010L0063).

### Cells

C57BL/6 mouse B16F10 (RRID:CVCL_0159) skin melanoma, PANO2 (RRID:CVCL_D627) pancreatic and Lewis lung carcinomas (LLC, RRID:CVCL_4358), as well as A549 (RRID:CVCL_0023) LUAD cells were from the National Cancer Institute Tumor Repository (Frederick, MD); human HEK293T (RRID:CVCL_0063) embryonic kidney cells were from the American Type Culture Collection (Manassas, VA); C57BL/6 mouse MC38 (RRID:CVCL_B288) colon adenocarcinoma cells were a gift from Dr Barbara Fingleton (Vanderbilt University, Nashville, TN, USA) (41, 42). B16F10 p*Kras*^G12C^, PANO2 p*Kras*^G12C^ and HEK293T p*Kras*^G12C^ cells were generated after stable transfection of B16F10, PANO2 and HEK293T cells with plasmids eGFP.KRASG12C-2B.retro.puro (Addgene ID 64372, RRID:Addgene_64372), eGFP.KRASG12C-2A.retro.puro (Addgene ID64373, RRID:Addgene_64373) and eGFP.KRASG12C-2B.retro.puro (Addgene ID 64372, RRID:Addgene_64372) respectively as described elsewhere (3). B16F10 p*C* and PANO2 p*C* were stably transfected with control (empty) overexpression vector eGFP.retro.puro (Addgene ID 64336, RRID:Addgene_64336) (3). HEK293T p*Kras*^G12C^ were stably transfected with p*LUC* (Cag.Luc.puro, Addgene ID 74409, RRID:Addgene_74409) or with plasmid enconding *Creb1* (p*Creb1*, Addgene ID 154942, RRID:Addgene_154942) constructed after *Creb1* cDNA introduction into a pCMVβ vector (Clontech, CA, #631719). Primary LUAD cells from C57BL/6 and FVB mice as well as *Trp53*- conditional LUAD cells were generated as described elsewhere (3, 27, 43) and *Creb1*^F/F^ LUAD cells were generated similarly. Briefly, C57BL/6 and *Trp53*^F/F^ (B6.129P2-*Trp53*^*tm1Brn*^/J #008462, RRID:IMSR_JAX:008462) mice received ten and FVB and *Creb1*^F/F^ mice on the FVB strain received four consecutive weekly intraperitoneal injections of urethane (1 g/kg) starting at 6 weeks of age and were killed ten months later. Lung tumors were isolated under sterile conditions, strained to single cell suspensions, and cultured for more than100 passages over two years to successfully yield tumor cell lines. Primary LUAD cells from FVB mice underwent RNA interference with the following lentiviral shRNA pools obtained from Santa Cruz Biotechnology (Palo Alto, CA): random control shRNA (sh*C*, sc-108080), and anti-Kras.shRNA (sh*Kras*, sc-43876) as described elsewhere (3). *Trp53*- conditional LUAD cells were stably transfected with vectors p*LUC* (Cag.Luc.puro, Addgene ID 74409, RRID:Addgene_74409) or with vectors encoding CRE recombinase p*CRE* (pPy-CAG-Cre::ERT2-IRES-BSD, Addgene ID 48760, RRID:Addgene_48760) (43, 44). *Creb1*^F/F^LUAD cells were stably transfected with vectors p*eGFP*, (retro-gfp-puro vector, Addgene ID 58249, RRID:Addgene_58249) (3), p*LUC* (Cag.Luc.puro, Addgene ID 74409, RRID:Addgene_74409) or with vectors encoding CRE recombinase (p*CRE*) (pPy-CAG-Cre::ERT2-IRES-BSD, Addgene ID 48760, RRID:Addgene_48760) (43, 44). For stable transfections, cells were transfected with 5 μg DNA using calcium phosphate. Stable clones were selected with antibiotic selection. All cell lines were cultured at 37°C in 5% CO_2_-95% air using full culture medium (DMEM supplemented with 10% FBS, 2 mM L-glutamine, 1 mM pyruvate, 100 U/ml penicillin, and 100 mg/ml streptomycin). For *in vivo* injections, cells were collected with trypsin, incubated with Trypan blue, counted by microscopy in a haemocytometer, their concentration was adjusted in PBS, and were injected in the skin, as described elsewhere (42, 43). Only 95% viable cells were used for *in vivo* injections. Cell lines were tested biannually for identity and stability by short tandem repeats, Sanger sequencing for driver mutations, and microarray and for *Mycoplasma Spp*. by PCR using designated primers.

### Mouse models

For the carcinogen-induced lung tumor model, six-week-old mice on the FVB background received one intraperitoneal urethane injection (1 g/Kg in 100 mL saline) and were sacrificed 6–7 months later (12, 18, 27). *Creb1*^F/F^ mice were bred > F12 to the susceptible FVB background (12, 18, 27) and six-week-old FVB.*Creb1*^F/F^ mice received 5 × 10^9^ intratracheal PFU Ad-CRE to achieve maximal recombination of the respiratory epithelium (12), were exposed to one intraperitoneal urethane injection (1 g/Kg in 100 mL saline) two weeks post Ad-CRE treatment and were sacrificed 6 months post-urethane treatment (12, 18, 27). Scgb1a1.CRE mice were bred > F12 to the FVB background and were then intercrossed with FVB.*Creb1*^F/F^ mice. Six-week-old FVB.*Creb1*^WT/WT^;Scgb1a1.CRE and FVB.*Creb1*^F/F^;Scgb1a1.CRE mice received one intraperitoneal urethane injection (1 g/Kg in 100 mL saline) and were sacrificed 6 months later. For mutant *KRAS*^G12D^-driven LUAD, C57BL/6 mice heterozygous for the loxP-STOP-loxP.*KRAS*^G12D^ transgene, LSL.*KRAS*^G12D^ mice, which express mutant *KRAS*^G12D^ in all somatic cells upon CRE-mediated recombination, received 5 × 10^8^ intratracheal PFU Ad-CRE and were killed after four months (12). LSL.*KRAS*^G12D^ mice were intercrossed with *Creb1*^F/F^ mice and B6.*Creb1*^F/F^;LSL.*KRAS*^G12D^ mice received 5 × 10^8^ intratracheal PFU Ad-CRE and were killed after four months (12). Control mice were a mixture of littermates negative for the transgenes of interest, including FVB.*Creb1*^WT/WT^ mice as appropriate controls for FVB.*Creb1*^F/F^ mice, FVB.*Creb1*^WT/WT^;Scgb1a1.CRE mice as appropriate controls for FVB.*Creb1*^F/F^;Scgb1a1.CRE mice, C57BL/6 WT mice as appropriate controls for LSL.*KRAS*^G12D^ mice and B6.*Creb1*^WT/WT^;LSL.*KRAS*^G12D^ mice as appropriate controls for B6.*Creb1*^F/F^;LSL.*KRAS*^G12D^ mice. For flank tumor formation, mice were anesthetized using isoflurane inhalation and received subcutaneous injections of 100 μL PBS containing the cells of interest into the rear flank. Three vertical tumor dimensions (δ1, δ2, and δ3) were monitored weekly and tumor volume was calculated using the formula π * δ1 * δ2 * δ3 / 6 as described elsewhere (42, 43). FVB mice and *Cxcr1*^*-/-*^ mice that were bred > F12 to the FVB background received 2 * 10^6^ *Creb1*^F/F^ LUAD cells stably transfected with p*CRE* or p*LUC. NOD/SCID* mice received 2 * 10^6^ HEK293T p*Kras*^G12C^ cells. C57BL/6, *Cxcr1*^*-/-*^ and *Cxcr2*^*+/-*^ mice received 10^6^ LLC or 10^6^ LUAD cells. C57BL/6 and *Cxcr1*^*-/-*^ mice received 10^6^ PANO2 or 10^6^ PANO2 p*Kras*^G12C^ cells. FVB mice received 10^6^ LUAD cells, followed by KG501 or ICG-001 treatments. For the treatment with the inhibitors, we intraperitoneally treated mice daily with KG501 (Sigma, St. Louis, MO, 70485) (15 mg/kg) for days 10-20 or 20-30 or with ICG-001 (AbMole, M2008) (5 mg/kg) for days 10-30 post-subcutaneous LUAD cells injection.

### Structural assessments in murine lungs

Mouse lungs were recoded (blinded) by laboratory members not participating in these studies and were always examined by two independent blinded participants of this study. The results obtained by each investigator were compared, and lungs were reevaluated if deviant by > 20%. Lungs and lung tumors were initially inspected macroscopically under a Stemi DV4 stereoscope equipped with a micrometric scale incorporated into one eyepiece and an AxiocamERc 5s camera (Zeiss, Jena, Germany) in transillumination mode, allowing for visualization of both superficial and deeply-located lung tumors. Tumor location was charted and diameter (δ) was measured. Tumor number (multiplicity) per mouse was counted and mean tumor diameter per mouse was calculated as the average of individual diameters of all tumors found in a given mouse lung. Individual tumor volume was calculated as πδ^3^/6. Mean tumor volume per mouse was calculated as the average of individual volumes of all tumors found in a given mouse lung, and total lung tumor burden per mouse as their sum. Lung volume was measured by saline immersion, and lungs were embedded in paraffin, randomly sampled by cutting 5 μm-thick lung sections (*n* = 10/lung), mounted on glass slides, and stained with hematoxylin and eosin for morphometry and histologic typing of lung tumors. For this, a digital grid of 100 intersections of vertical lines (points) was superimposed on multiple digital images of all lung sections from lung tissue of a given mouse using Fiji academic freeware (RRID:SCR_002285) (45). Total lung tumor burden was determined by point counting of the ratio of the area occupied by neoplastic lesions versus total lung area and by extrapolating the average ratio per mouse to total lung volume (46) .The results of this stereologic approach were compared with the macroscopic method detailed above and were scrutinized if deviant by > 20%.

### Histology

Lungs were fixed with 10% formalin overnight and were embedded in paraffin. Cells were grown on glass coverslips and fixed with 100% methanol at 4 °C for 5 minutes. Five-μm-thick tissue sections were counterstained with hematoxylin and eosin (Sigma, St. Louis, MO) or, similarly to the cells coverslips, were incubated with the indicated primary antibodies (Reagents) overnight at 4 °C followed by Envision/diaminobenzidine detection (Dako, Glostrup, Denmark) and hematoxylin counterstaining/mounting (Entellan; Merck Millipore, Darmstadt, Germany). For isotype control, primary antibody was omitted. Bright- field images were captured with an AxioLab.A1 microscope connected to an AxioCamERc 5s camera (Zeiss, Jena, Germany). Digital images were processed with Fiji. P-CREB nuclear and CXCR1cytoplasmic immunostaining intensity of the lung tissues with regard to staining intensity and fraction of stained cells was defined in ten independent lung sections by visual analog scale. P-CREB nuclear immunostaining intensity of the cells was assessed as percentage of P-CREB+ cells (%) in four independent fields. Staining was evaluated by two blinded readers. Multi-color immunohistochemistry was conducted on a cohort of early-stage non-small cell lung cancer patients as described elsewhere (https://jitc.bmj.com/content/9/2/e001469) with polyclonal rabbit anti-Neutrophile Elastase antibody (Abcam, Cambridge, UK; ab68672, RRID:AB_1658868) diluted 1:100 and monoclonal mouse anti-pan Cytokeratin antibody (AE1/AE3, Zytomed Systems, Berlin, Germany) diluted 1:300. Neutrophil elastase (NE) was detected using OPAL 570 and pan-CK was detected using OPAL 690 fluorophores both at 1:150 dilution and 10 min OPAL-TSA reaction time. Heat-induced antigen retrieval (HIER) was conducted using an inverter microwave with 1 min at 1000W and 10 min at 100W corresponding to one cycle. NE was placed at 1st cycle and pan-CK at 2nd cycle in the mIF panel. Slides were scanned at a Vectra Polaris 1.0 fluorescent slide scanner (Akoya Biosciences, Marlborough, MS, USA) at 20× (0.5 μm/pixel) magnification using the following filters and exposure times: DAPI (1.81 ms), OPAL 570 (16 ms), OPAL 690 (13 ms) and Sample AF (100 ms). Phenochart 1.1 software (Akoya Biosciences, RRID:SCR_019156) was used for image annotation and grid placement for tissue microarray (TMA) analysis. Images were analyzed using inForm 2.5 software (Akoya Biosciences, RRID:SCR_019155). Image analysis was conducted with representative images sampled from across all 5 TMAs to set up an algorithm for batch analysis. Tissue segmentation was performed based on the pan- Cytokeratin (CK) signal to discriminate tumor from stromal areas as well as tissue-free regions. Adaptive cell segmentation was performed and two cell classifiers trained to identify either, NE+ cells vs NE- cells or panCK+ cells vs panCK- cells. Both algorithms were used to batch analyze all images. Results from both cell classifiers were consolidated into one file using phenoptr 0.2.4. (http://akoyabio.github.io/phenoptr) and phenoptrReports 0.2.5. (http://akoyabio.github.io/phenoptrReports) R packages and NE+ cells exported as cell density normalized to the respective tissue segmentation result’
ss area (stroma/ tumor) and expressed as NE+ cells/mm2. Since the TMAs harbored ≥ 2 punches per patient, the mean was calculated from individual punches and used for downstream analysis.

### Bronchoalveolar lavage (BAL)

BAL was performed using three sequential aliquots of 1 mL sterile ice-cold PBS. Fluid was combined and centrifuged at 300 g for 10 min at 4 °C to separate cells from supernatant. The cell pellet was resuspended in 1 ml PBS containing 2% FBS and 0.1% azide, and the total cell count was determined using a grid hemocytometer according to the Neubauer method. Neutrophil content was determined by counting 400 cells on May-Grünwald-Giemsa-stained cytocentrifugal specimens as well as by determining numbers of CD45^+^ CD11b^+^ Gr1^+^ CXCR1^+^ cells by flow cytometry. For flow cytometry, the antibodies FITC-labeled rat monoclonal anti-mouse CD45 (clone 30-F11, eBioscience; 11- 0451-82, RRID:AB_465050), PE-labeled rat monoclonal anti-mouse CD11b (clone M1/70, eBioscience; 12-0112-82, RRID:AB_2734869), PE-Cyanine7-labeled rat monoclonal anti-mouse Gr1 (clone RB6-8C5, eBioscience; 25-5931-82, RRID:AB_469663), and mouse monoclonal anti-mouse CXCR1 (Abcam, Cambridge, UK; ab10400, RRID:AB_297141) were used in a concentration of 1μg/μl.

### Cellular Assays

*In vitro* cancer cell proliferation was determined using the MTT method. Briefly, on day 0, 150 μL of a 2 × 10^4^ cell suspension (in quadruplicates) were plated in four 96-well plate wells. Each day, (for the following four days), 15 μL of MTT working solution (5 mM MTT in PBS) was added to each cell culture well of a single plate. The plate was left for 4 h at 37 °C in a 5% CO_2_ humidified incubator followed by the addition of 100 μL acidified isopropanol and subsequent sediment solubilization. Absorbance was measured in a MR-96A microplate reader (Mindray, Shenzhen, China) at 492 nm. Cytoplasmic extracts from *Creb1*^F/F^ LUAD cells, transfected with p*LUC* or p*CRE*, were assayed for CCL2 (PeproTech; 900-K126) and CXCL5 (Sigma, St. Louis, MO, RAB0131) activity using commercially available ELISA kits.

### Immunoblotting

Nuclear and cytoplasmic protein extracts from LLC, A549, LUAD cells from urethane-treated FVB mice, *Creb1*^F/F^ LUAD cells transfected with p*LUC* or p*CRE*, HEK293T p*LUC*, HEK293T p*Kras*^G12C^, *Trp53*- conditional LUAD cells transfected with p*LUC* or p*CRE*, LUADsh*C*, LUADsh*Kras*, B16F10 p*C*, B16F10 p*Kras*^G12C^, PANO2 p*C*, and PANO2 p*Kras*^G12C^ were prepared using NE-PER extraction kit (Thermo, Waltham, MA), were separated by 12% sodium SDS-PAGE, and were electroblotted to PVDF membranes (Merck Millipore, Darmstadt, Germany). Membranes were probed with specific antibodies (Reagents), and were visualized by film exposure after incubation with enhanced chemiluminescence substrate (Merck Millipore, Darmstadt, Germany).

### Genomic studies and transcriptome analyses

Triplicate cultures of *Creb1*^F/F^ LUAD cells transfected with p*LUC* or p*CRE* were subjected to RNA extraction using Trizol (Invitrogen, Carlsbad, CA) followed by column purification and DNA removal (Qiagen, Hilden, Germany) and reverse transcription using Superscript III (Invitrogen, Thermo Fisher Scientific, Waltham, MA). RT-PCR was performed using first strand synthesis with specific primers (Reagents) and SYBR FAST qPCR Kit (Kapa Biosystems, Wilmington, MA) in a StepOne cycler (Applied Biosystems, Carlsbad, CA). Ct values from triplicate reactions were analyzed with the 2^-ΔCT^ method (47) relative to glycuronidase beta (*Gusb*). For microarray, 5 μg pooled RNA was quality tested on an ABI2000 bioanalyzer (Agilent Technologies, Sta. Clara, CA), labeled, and hybridized to GeneChip Mouse Gene 2.0 ST arrays (Affymetrix, Sta. Clara, CA). All data were deposited at GEO (http://www.ncbi.nlm.nih.gov/geo/; Accession ID: GSE156513), were subjected to WikiPathway analysis (48) and were analyzed on the Gene Expression and Transcriptome Analysis Consoles (Affymetrix, Santa Clara, CA, RRID:SCR_018718) using as cut-off differential gene expression > 2. Murine CREB signatures were compared with normal tissue, smoker LUAD and non-smoker LUAD microarray data (GEO dataset GSE43458; https://www.ncbi.nlm.nih.gov/geo/query/acc.cgi?acc=GSE43458, RRID:SCR_005012) (49-51). Humanized *CREB1* signatures were derived from murine *Creb1* signatures using http://www.ensembl.org/biomart/martview. Hierarchical clustering of BATTLE study patients by the *CREB1* signature was performed using GEO series GSE43458. Human LUAD patient survival analyses were done using Kaplan–Meier Plotter (http://kmplot.com/analysis/index.php?p=service&cancer=lung, RRID:SCR_018753) (52) and parameters auto-select best cutoff, compute median survival, censor at threshold, and histologic subtype LUAD and LUSC. GSEA was performed with the Broad Institute pre-ranked GSEA module (https://www.gsea-msigdb.org/gsea/index.jsp, RRID:SCR_003199) (53) using BATTLE study transcriptomes from GEO series GSE43458. GEO dataset GSE43458 (https://www.ncbi.nlm.nih.gov/geo/query/acc.cgi?acc=GSE43458) (49) and GEO dataset GSE31852 (https://www.ncbi.nlm.nih.gov/geo/query/acc.cgi?acc=GSE31852) (50,51) were combined and the transcription levels of *CREB* were assessed in the corresponding LUAD tissues and juxtatumoral lungs compared with housekeeping gene *ACTB. Kras* sequencing is described elsewhere (3). Briefly, one μg RNA was reverse-transcribed using Oligo(dT)_18_ primer and Superscript III (Invitrogen, Thermo Fisher Scientific, Waltham, MA). *Kras* cDNA was amplified in PCR reaction using the corresponding primers (Reagents) and Phusion Hot Start Flex polymerase (New England Biolabs, Ipswich, MA). cDNA fragments were purified with NucleoSpin gel and PCR clean-up columns (Macherey-Nagel, Düren, Germany) and were Sanger- sequenced using their corresponding primers by VBC Biotech (Vienna, Austria).

### ENCODE transcription factor analyses

ChIP-seq datasets from the ENCODE Transcription Factor Targets dataset (RRID:SCR_006793) and from the CHEA Transcription Factor Targets dataset (RRID:SCR_005403) were used. The CREB binding sequence motif from the ENCODE portal (https://www.encodeproject.org/) with the identifier: ENCFF576PUH was downloaded. Motif-based sequence analysis using T-Gene tool of MEME suite 5.3.0 (RRID:SCR_001783) was deployed in order to predict regulatory links between loci of CREB binding and the target genes (annotation with reference genome Homo sapiens hg38, UCSC). *e* represents the statistical significance of the motif in terms of probability to be found in similarly sized set of random sequences.

### Kaplan-Meier plotter analysis

For these analyses performed at Kaplan–Meier Plotter (http://kmplot.com/analysis/index.php?p=service&cancer=lung, RRID:SCR_018753) (52) and parameters auto-select best cutoff, compute median survival, censor at threshold, and histologic subtype LUAD and LUSC, eight human orthologues of 15 top CREB-induced genes were non inverted: ERO1L (218498_s_at), CIP1 (202284_s_at), ACAN (207692_s_at), PLCE1 (205111_s_at), HEXA (201765_s_at), SLCO2A1 (204368_at), MT2A (212185_x_at), CLDN9 (214635_at). Twelve human orthologues of 15 top CREB- suppressed genes were inverted: IL1A (210118_s_at), AQP8 (206784_at), CD93 (202878_s_at), ARHGAP18 (225166_at), CXCL6 (206336_at), DOCK11 (226875_at), RET (211421_s_at), SCYB1 (204470_at), LY6G6C (207114_at), PPBP (214146_s_at), TBC1D15 (230072_at), TRAC (209670_at).

### Human protein atlas analyses

The immunoreactivity of CREB and CXCR1 in the tumor site and the stroma of 43 and 22, respectively, human LUAD tissues available from The Human Protein Atlas (http://www.proteinatlas.org) (53) was assessed by visual analog scale scoring. Representative images (https://images.proteinatlas.org/19150/44741_B_3_4.jpg and https://images.proteinatlas.org/31991/135678_B_1_4.jpg) were obtained from https://www.proteinatlas.org/ENSG00000118260-CREB1/pathology/lung+cancer#img and https://www.proteinatlas.org/ENSG00000163464-CXCR1/pathology/lung+cancer#img respectively, available from v19.proteinatlas.org.

### Statistics

Sample size was calculated using G*power (http://www.gpower.hhu.de, RRID:SCR_013726) assuming *α* = 0.05, *β* = 0.05, and effect size *d* = 1.5. No data were excluded. Animals were allocated to treatments by alternation and transgenic animals were enrolled case-control-wise. Data acquisition was blinded on samples previously coded by a non-blinded investigator. All data were examined for normality by Kolmogorov-Smirnov test. Data are given as raw data points (circles), rotated kernel density plots (violins), medians (dashed lines), and interquartile ranges (dotted lines), mean ± SD, or median ± interquartile range, as indicated and as appropriate. Sample size (*n*) refers to biological replicates. Differences in means were examined by Student’s t-test, one-way, or two-way ANOVA with Bonferroni post- tests, as appropriate, and in medians by Mann-Whitney U-test or Kruskal-Wallis test with Dunn’s post-hoc tests. Hypergeometric tests were performed according to the Graeber Lab at https://systems.crump.ucla.edu/hypergeometric/index.php. Probability (*P*) values are two-tailed and *P* < 0.05 was considered significant. Analyses and plots were done on Prism v8.0 (GraphPad, La Jolla, CA, RRID:SCR_002798).

## AUTHOR CONTRIBUTIONS

GAG, GN, AM, and IL performed *in vitro* experiments, *in vivo* experiments, histology, microscopy, GSEA, and wrote the draft of the manuscript; FK, ET, NS, IG, MS, and MI performed flow cytometry, pharmacologic CREB inhibition, *in vivo* experiments, genotyping, cell culture, RNA isolation, tumor cell and carcinogen injections, flow cytometry, and tissue processing; AK analyzed human microarray data; TM provided critical analytical tools; CR, TG, and SM performed multi-color immunohistochemistry and analyses: GTS designed, funded, and guided the study, analyzed the data, wrote the final version of the manuscript, and is the guarantor of the study’s integrity. All authors critically reviewed and edited the paper for important intellectual content and approved the final submitted version.

## ACKNOWLEDGEMENTS

This work was supported by European Research Council grants #260524 and #679345, the Graduate College GRK2338 of the German Research Society (DFG), the target validation project for pharmaceutical development ALTERNATIVE of the German Ministry for Education and Research (BMBF), and the German Center for Lung Research (DZL) (to GTS); the Greek State Scholarship Foundation/European Union Social Fund/Greece program NSRF 2014-2020 and grant MIS-5033021 (to IG, MS, and IL); the General Secretariat for Research and Innovation and Hellenic Foundation for Research and Innovation grant #1853a (to MS); and a Hellenic Association for Molecular Cancer Research Award (to AM). The authors thank the BioMaterialBank North, funded by the Airway Research Center North, the DZL, popgen 2.0 network (P2N), and BMBF grant 01EY1103, for patient tissues.

## Notes

**Conflict of interest:** The authors have declared that no conflict of interest exists.

### Competing Interest Statement

The authors have declared no competing interest.

