## Supplemental Figures for "Immune-evasion of *KRAS*-mutant lung adenocarcinoma mediated by cAMP response element-binding protein"

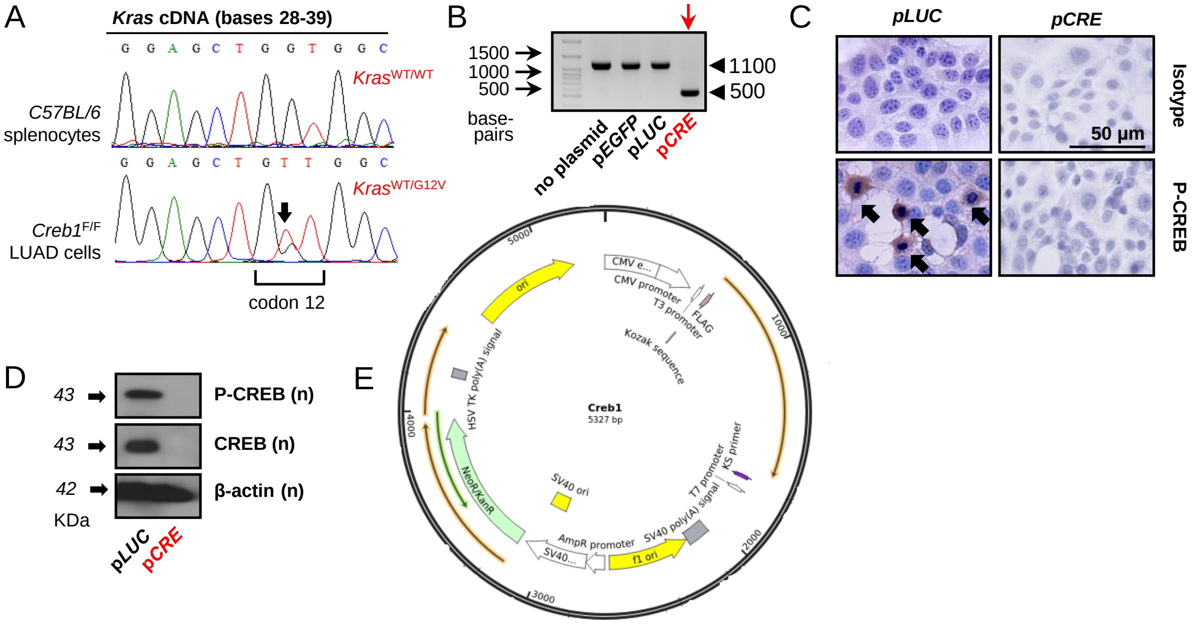


**Fig. S1. Validation of primary *Creb1*-conditional LUAD cells isolated from a urethane-treated *Creb1*$^{F/F}$ mouse. (A)** cDNA Sanger sequencing shows the presence of a heterozygous *Kras*$^{G12V}$ mutation (black arrow). **(B-D)**Validation of recombination resulting in *Creb1* deletion (red arrow) upon stable transfection with plasmid encoding CRE recombinase (p*CRE*) and of intact *Creb1* locus in primary cells stably transfected with no plasmid, plasmid encoding EGFP (p*EGFP*) or *Photinus pyralis* luciferase (p*LUC*). Shown are representative PCR (B), immunocytochemistry (C), and immunoblot (D) results. PCR bands at 500 and 1100 bp represent, respectively, the PCR products of recombined conditional (deleted) and non-recombined conditional *Creb1* transgene loci. **(E)** Creb1 construct for eukaryotic overexpression.

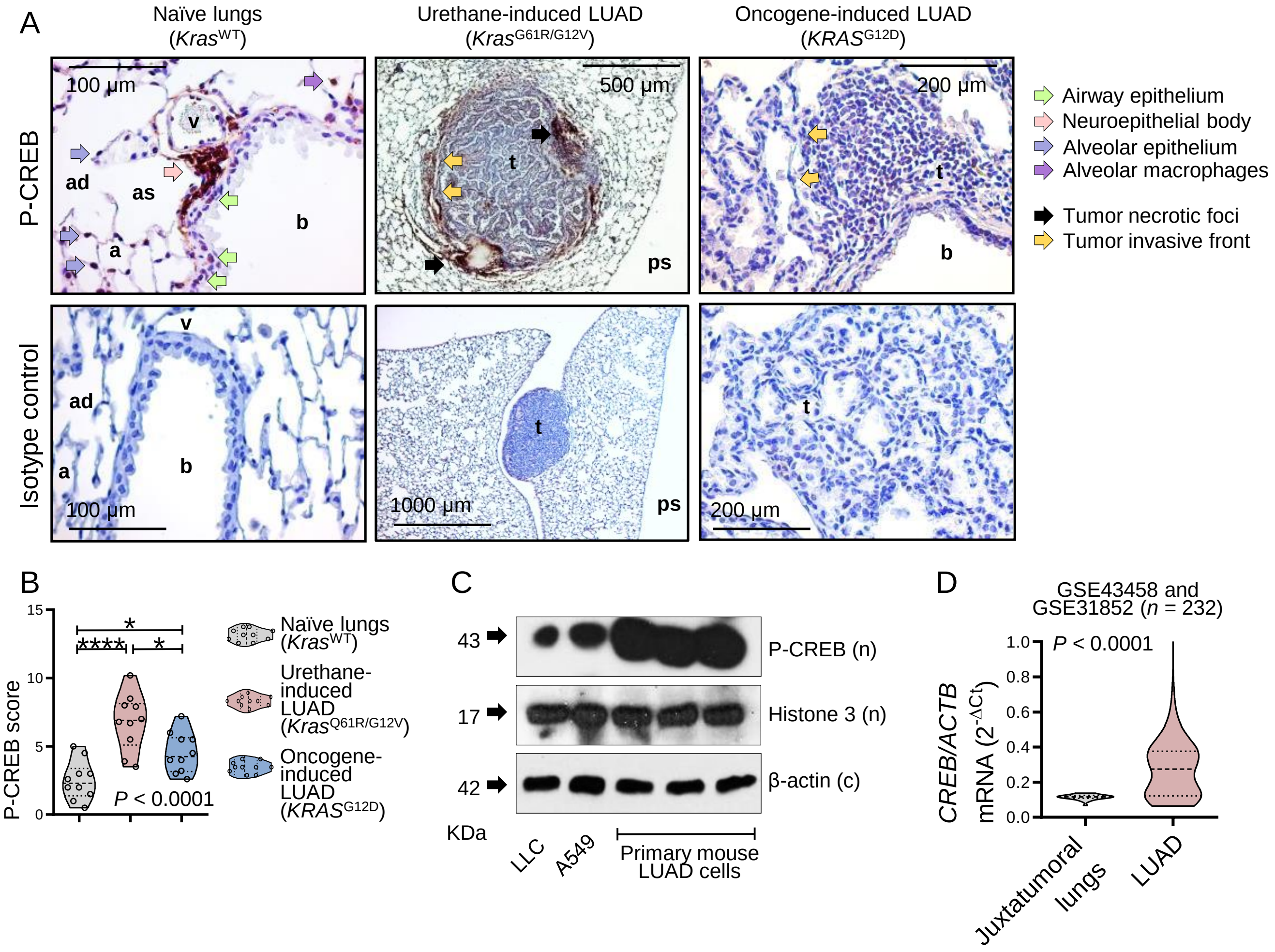


**Fig. S2. CREB is overexpressed in *KRAS*-mutant mouse and human LUAD. (A)** Representative images of P-CREB (brown) and hematoxylin (blue)-stained lung sections from naïve FVB mice (left), FVB mice at six months post-urethane (middle), and LSL.*KRAS*$^{G12D}$ mice at four months post-intratracheal Ad-CRE (right) ($n$ = 10/group). For isotype controls, the primary antibody was omitted. Note P-CREB expression across the respiratory epithelium, in neuroepithelial bodies, and alveolar macrophages. Note also increased P-CREB expression in lung tumors, especially around necrotic foci and adjacent to invasive fronts. a, alveolus; ad, alveolar duct; as, alveolar sac; b, bronchus; ps, pleural space; t, tumor; v, vein. **(B)** Data summary ($n$ = 10 mice/group) from (A) shown as raw data points (circles), rotated kernel density plots (violins), medians (dashed lines), and interquartile ranges (dotted lines). $P$, probability, one-way ANOVA; * and ****: $P < 0.05$ and $P < 0.0001$, respectively, for the indicated comparisons, Tukey's post-tests. **(C)** Immunoblots of nuclear (n) and cytoplasmic (c) protein extracts of three different primary urethane-induced LUAD cells (24) compared with mouse (LLC) and human (A549) LUAD cell lines show increased nuclear P-CREB in the former primary cells. **(D)** Average *CREB* expression normalized to *ACTB* in LUAD ($n$ = 202) and healthy lungs ($n$ = 30) from the indicated GEO datasets. Data shown as rotated kernel density plots (violins), medians (dashed lines), and interquartile ranges (dotted lines). $P$, probability, Mann-Whitney U test.

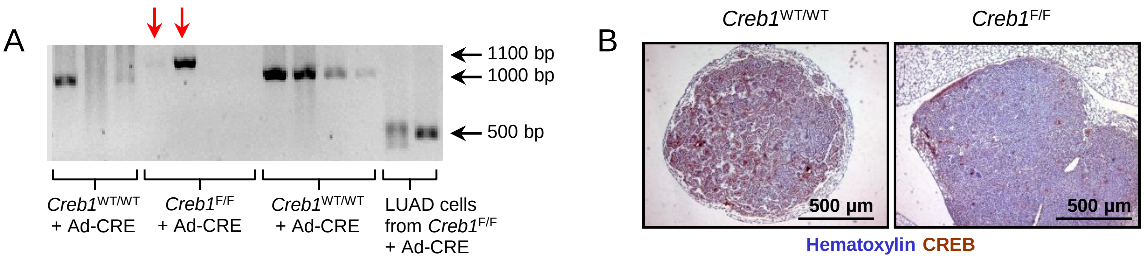


C

| ***Creb1* detection by PCR** | YES | NO | *P* |
|---|---|---|---|
| ***Creb*$^{WT/WT}$ Detected** | 22 | 3 | 1.0000 |
| **Unrecombined *Creb*$^{F/F}$ detected** | 14 | 1 | |

D

| **P-CREB immunoreactivity** | YES | NO | *P* |
|---|---|---|---|
| ***Creb*$^{WT/WT}$** | 48 | 0 | 0.3333 |
| ***Creb*$^{F/F}$** | 23 | 1 | |

**Fig. S3. CREB is required for urethane-induced *KRAS*-mutant LUAD.** *Creb1*$^{WT/WT}$ and *Creb1*$^{F/F}$ mice on the FVB strain received intratracheal Ad-CRE (5 x $10^9$ PFU) followed by intraperitoneal 1 g/Kg urethane after 2 weeks. Lung tumors were harvested 6 months after urethane. **(A, B)** Representative images of PCR of lung tumor DNA targeted at the recombination event of the *Creb1* transgene (A) and of P-CREB (brown) and hematoxylin (blue)-stained lung tumor sections (B). PCR bands at 500, 1000, and 1100 bp represent, respectively, the PCR products of recombined conditional (deleted), wild-type, and non-recombined conditional *Creb1* transgene loci. Note also the presence of P-CREB immunoreactivity of lung tumors from *Creb1*$^{F/F}$ mice. **(C, D)** Pooled results from analyses as in (A, B) indicate that the few tumors observed in *Creb1*$^{F/F}$ mice result from the minority of non-recombined (i.e., *Creb1*-deleted) respiratory epithelium, which represent approximately 10% of all respiratory epithelial cells in this model, as shown previously (12). Shown are number (*n*) of lung tumors with and without *Creb1* deletion (C) and P-CREB immunoreactivity (D). *P*, probability, Fisher's exact test.

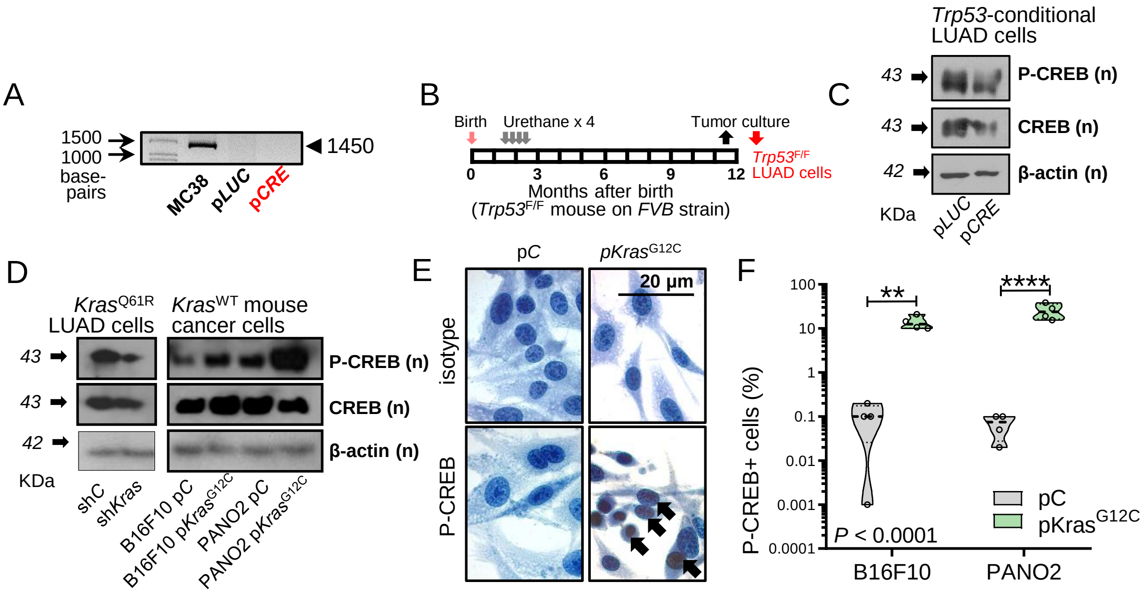


**Fig. S4. CREB activation is driven by mutant *Kras* and not *Trp53*. (A)** *Trp53* PCR of MC38 murine colon adenocarcinoma cells (*Trp53*$^{R178P}$ mutant) and *Creb1*$^{F/F}$ LUAD cells expressing p*LUC* and p*CRE* indicates *Trp53* deletion of LUAD cells. PCR band at 1450 bp represents the PCR product of the WT *Trp53* locus. **(B, C)** *Trp53*$^{F/F}$ LUAD cells were derived from a LUAD induced by four weekly i.p. injections of 1 g/Kg urethane in a FVB *Trp53*$^{F/F}$ mouse, were stably transfected with p*LUC* or p*CRE*, and were assessed for CREB activation by immunoblots (n, nuclear; c, cytoplasmic). **(D-F)** Representative immunoblots (D), immunocytochemistry images (E), and immunocytochemistry data summary (F) of *Kras*$^{Q61R}$ LUAD cells stably transfected with control shRNA (sh*C*) or anti-*Kras*-specific shRNA (sh*Kras*) and of *Kras*$^{WT}$ murine skin melanoma (B16F10) and pancreatic adenocarcinoma (PANO2) cells stably transfected with a control vector (p*C*) or mutant p*Kras*$^{G12C}$. D: n, nuclear; c, cytoplasmic. E: brown, P-CREB; blue, hematoxylin; arrows, nuclear P-CREB. F: Data summary ($n$ = 4 independent experiments/group) from (E) shown as raw data points (circles), rotated kernel density plots (violins), medians (dashed lines), and interquartile ranges (dotted lines). $P$, probability, two-way ANOVA; ** and ****: $P < 0.01$ and $P < 0.0001$, respectively, for the indicated comparisons, Bonferroni post-tests.

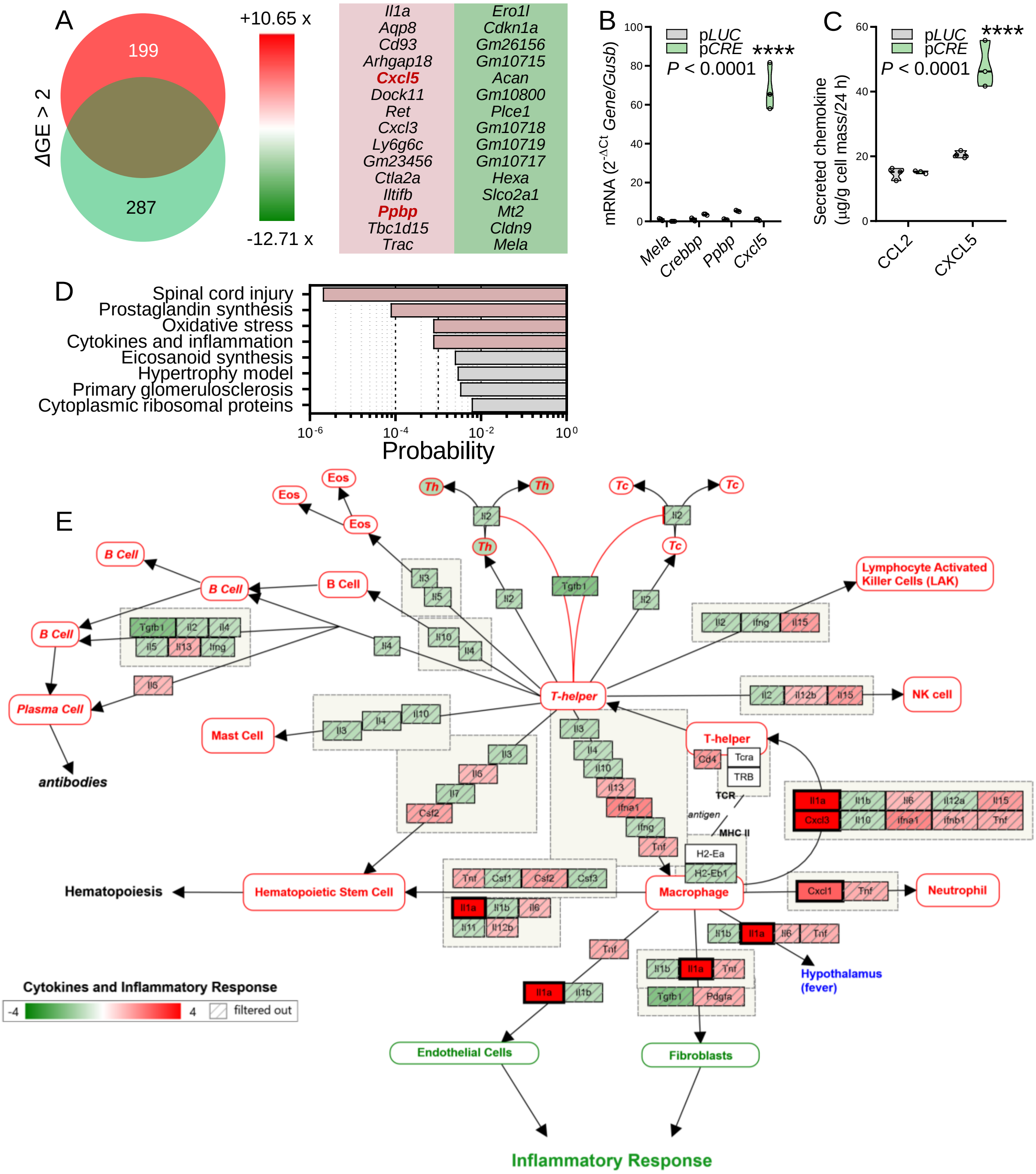


**Fig. S5. CREB suppresses CXCR1/2 ligand expression in the cellular context of mutant *Kras*. (A)** Venn diagram of transcriptome analysis representing genes responsive (*Δ*GE >2) to *Creb1* deletion. Red, induction; green, suppression; GE, gene expression. **(B)** *Mela*, *Crebbp*, *Ppbp*, and *Cxcl5* mRNA expression normalized to *Gusb* by RT-PCR and **(C)** CCL2 and CXCL5 protein secretion by ELISA of *Creb1*$^{F/F}$ LUAD cells transfected with p*LUC* ($n$ = 3) or p*CRE* ($n$ = 3). Data are shown as raw data points (circles), rotated kernel density plots (violins), medians (dashed lines), and interquartile ranges (dotted lines). $P$, probability, two-way ANOVA. ****: $P < 0.0001$ compared with p*LUC*, Sidak's post-tests. Only statistically significant differences are indicated. **(D)** WikiPathway analysis showing pathways significantly overrepresented in the *Creb1* signature. **(E)** Detailed schematic of perturbations induced by Creb1 deletion in the WikiPathway "Cytokines and inflammation".

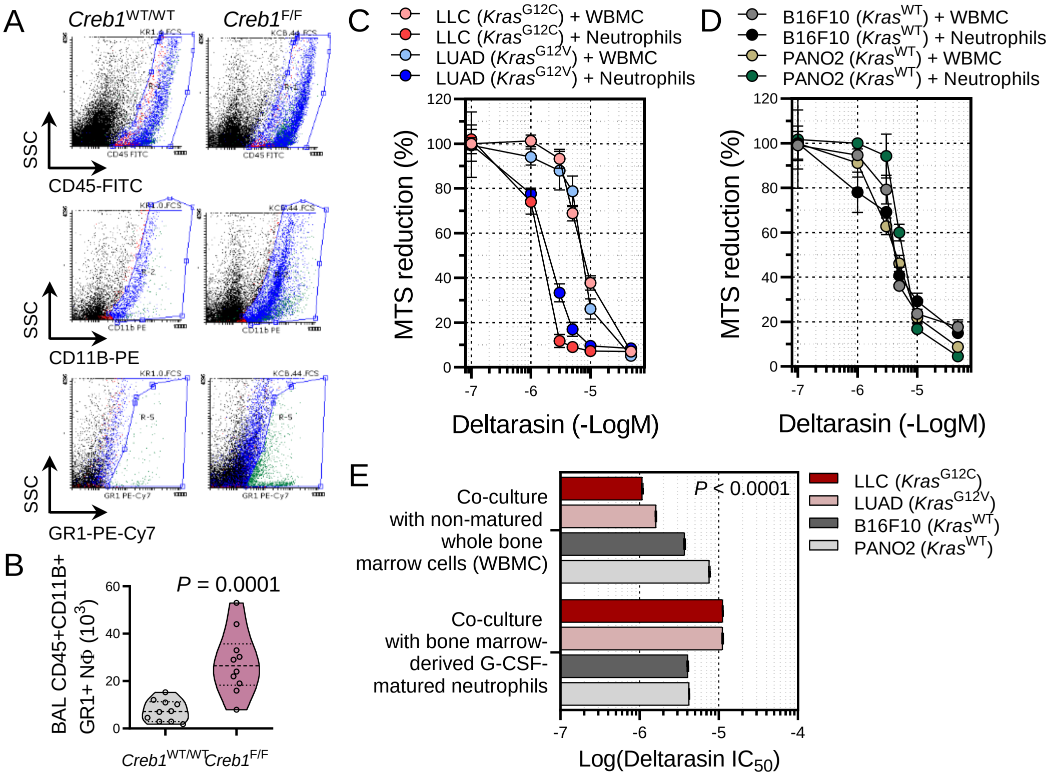


**Fig. S6. CREB signaling prevents neutrophil influx into the LUAD-affected lungs. (A)** Representative flow cytometric gating strategy of BAL from LUAD-bearing *Creb1*$^{WT/WT}$ and *Creb1*$^{F/F}$ mice. **(B)** Data summary ($n$ = 10/group) of flow cytometry for CD45+CD11b+Gr1+ cells from (A) shown as raw data points (circles), rotated kernel density plots (violins), medians (dashed lines), and interquartile ranges (dotted lines). *P*, probability, Mann-Whitney U-test. **(C-E)** Different murine cancer cell lines with mutant (C) or wild-type (D) Kras were incubated with DMEM + 50% conditioned media from whole bone marrow cells (WBMC) or from bone marrow-derived neutrophils (NΦ) generated after 2-day culture of WBMC with 20 ng/mL recombinant granulocyte colony stimulating factor (G-CSF). Cells were then treated with different concentrations of the KRAS inhibitor deltarasin. Shown in (C, D) are data summaries from three independent experiments as mean (circles), SD (bars) and in (E) is data summary ($n$ = 3) of deltarasin 50% inhibitory concentrations ($IC_{50}$) as mean (columns), SD (bars), and 2-way ANOVA probability (*P*).

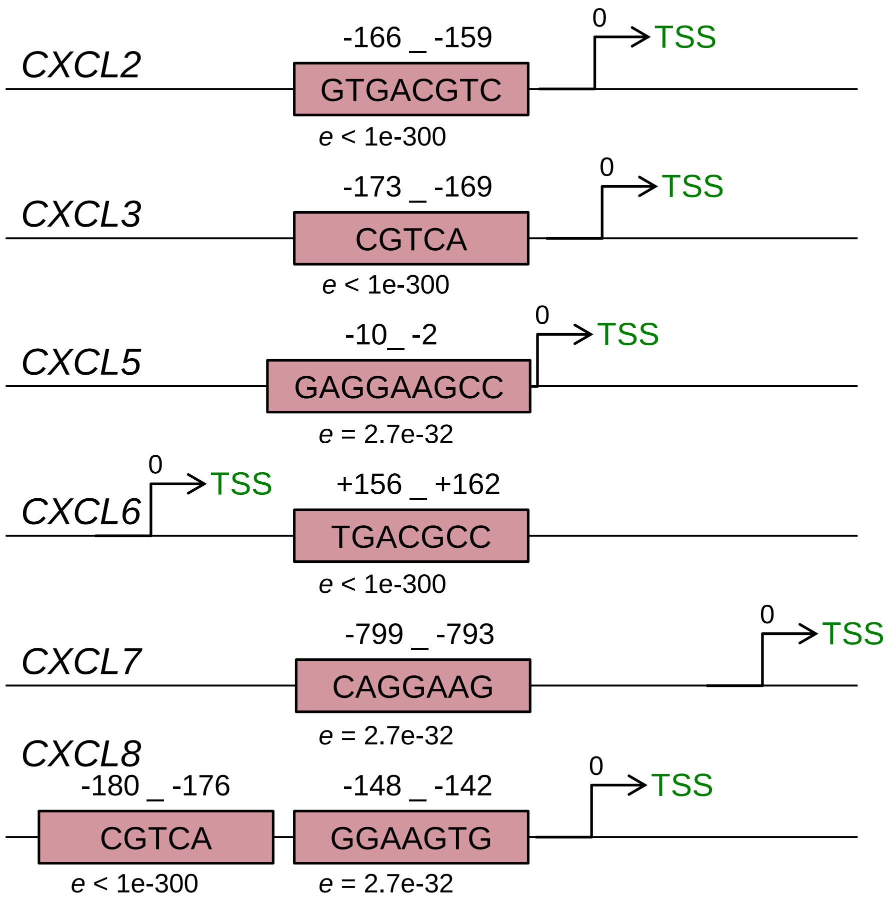


**Fig. S7. CREB regulates the transcription of CXCR1 ligands.** The CREB binding sequence motif was downloaded from the ENCODE portal (https://www.encodeproject.org/) with the identifier ENCFF576PUH. We employed the Motif-based sequence analysis T-Gene tool of MEME suite 5.3.0 in order to predict regulatory links between loci of CREB binding and the target genes [annotation with reference genome Homo sapiens hg38 (UCSC)]. *e* represents the statistical significance of the motif in terms of probability to be found in similarly sized set of random sequences. TSS: Transcription Start Site.

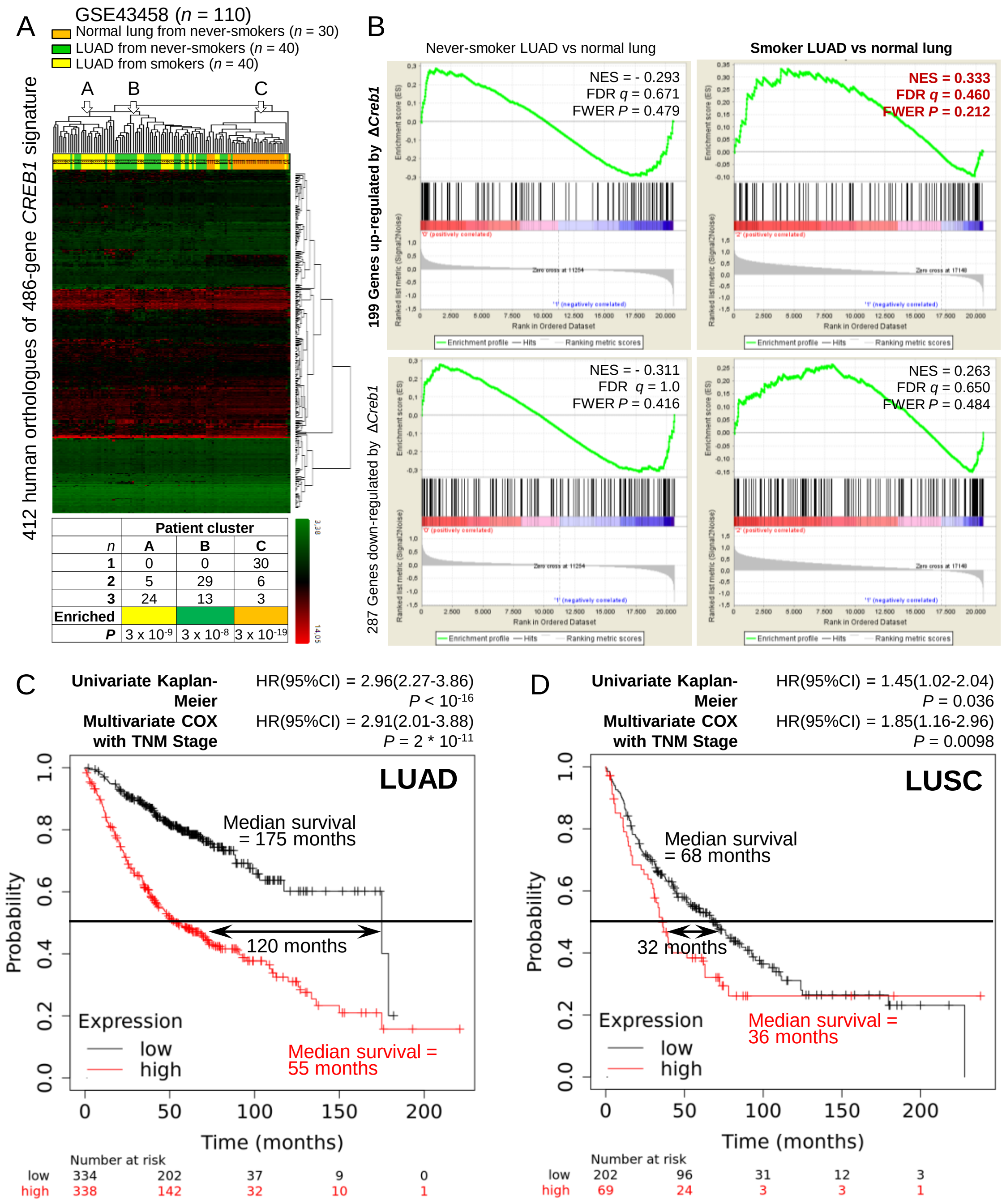


**Fig. S8. CREB1 signatures in human LUAD. (A, B)** Unsupervised hierarchical clustering (A) and pre-ranked gene set enrichment analysis (GSEA; B) of 30 normal lung tissues from never smokers (orange), 40 LUAD tissues from never smokers (green), and 40 LUAD tissues from smokers (yellow) from GEO dataset GSE43458 by the humanized *CREB1* signature derived from Supplemental Figure 6A. *P*, probability, hypergeometric test. Shown in (B) are enrichment plots, normalized enrichment scores (NES), false discovery rate *q* values (FDR), and family-wise error rates (FWER). Note significant (FDR $q < 0.5$) enrichment of the genes up-regulated by *Creb1* deletion in smokers' LUAD (red fonts). **(C, D)** Kaplan-Meier survival estimates of 672 patients with LUAD and 271 patients with squamous cell lung carcinoma (LUSC) from http://kmplot.com/ stratified by expression of the human orthologues of the 15 top genes induced (not inverted) or suppressed (inverted) by *Creb1* deletion from Supplemental Figure 6A. Shown are univariate and multivariate (COX regression by TNM stage) hazard ratios (HR) and probabilities (*P*).

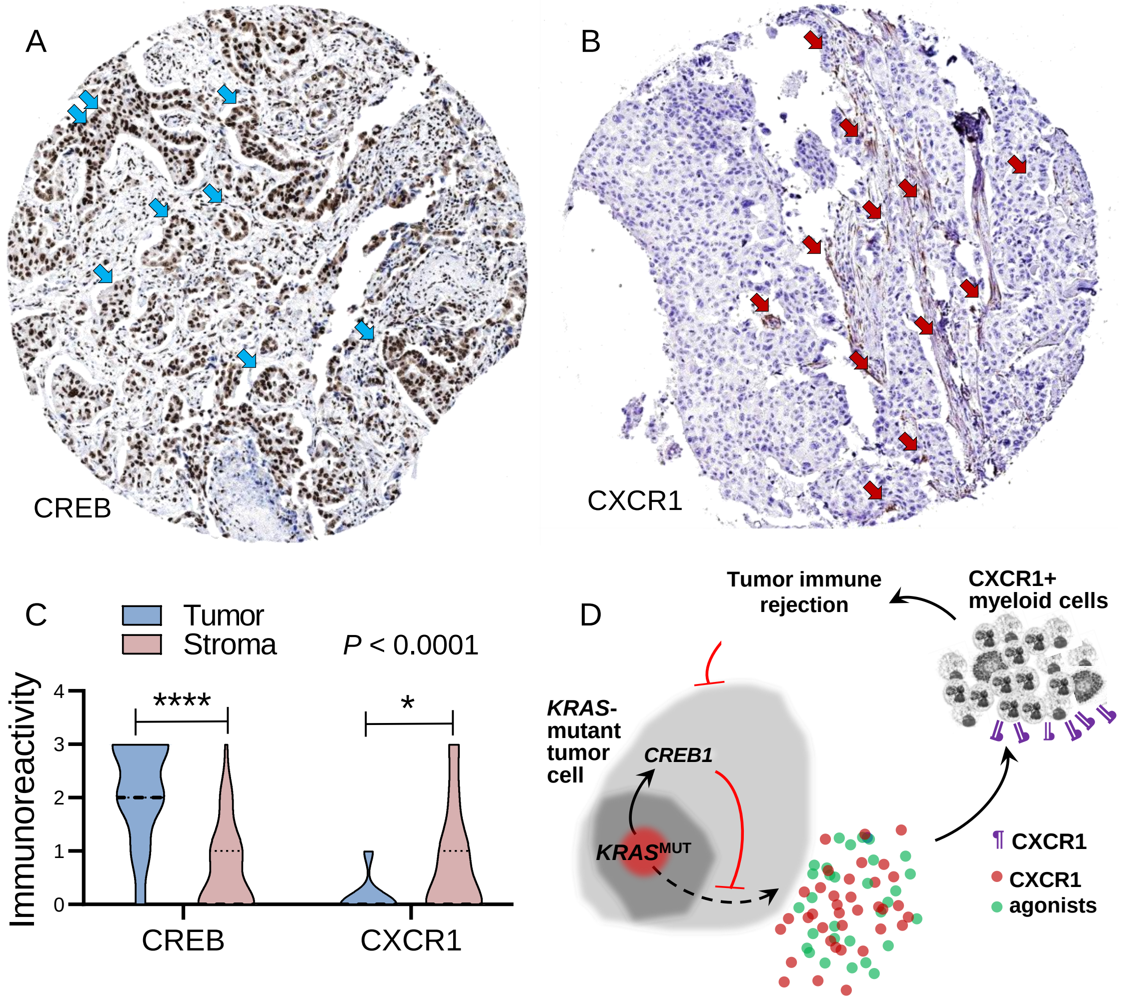


**Fig. S9. CREB and CXCR1 in LUAD from the human protein atlas. (A-C)** Representative images (A, B) and data summary (C) of CREB and CXCR1 (brown) and hematoxylin (blue)-stained sections (*n* = 43 and 22, respectively) of human LUAD tissues from the human protein atlas (http://www.proteinatlas.org). Note the predominant immunoreactivity of CREB in tumor cells (cyan arrows) and of CXCR1 in the tumor stroma (red arrows). Data summary (C) shows raw data points (circles), rotated kernel density plots (violins), medians (dashed lines), and interquartile ranges (dotted lines). *P*, probability, two-way ANOVA. * and ****: $P < 0.05$ and $P < 0.0001$, respectively, for the indicated comparisons, Bonferroni post-tests. **(D)** Schematic of proposed mechanism of CREB1-mediated immune evasion of KRAS-mutant LUAD: mutant KRAS activates CREB1, which functions to suppress CXCR1 agonist secretion and tumoricidal neutrophil recruitment.
